# Experiences and theory reveal antimicrobial blue light killing efficiency decrease as biofilm grows in a Pseudomonas aeruginosa model

**DOI:** 10.1101/2025.02.26.636652

**Authors:** Giacomo Insero, Nidia Maldonado-Carmona, Thomas Panier, Giovanni Romano, Nelly Henry

**Author notes:** Nelly Henry, Giovanni Romano. **Email:**. these authors contributed equally to the manuscript (Giacomo Insero, Nidia Maldonado-Carmona). current affiliation: University of Florence, Department of Experimental and Clinical Biomedical Sciences “Mario Serio”, Florence, Italy (Nidia Maldonado-Carmona).

## Abstract

Recently, the use of antimicrobial blue light (aBL) has gained interest across various applications. However, a comprehensive framework that addresses the key factors driving bacterial photoinhibition remains lacking—particularly concerning biofilms, the predominant bacterial lifestyle. The goal of this work was to evaluate the potential of photokilling in this wide-spread microbial adherent community type, and to decipher the specific mechanisms at stake. To investigate aBL killing efficiency, we conducted experiments in a *Pseudomonas aeruginosa* biofilm model using a well-defined millifluidic device that allows real-time microscopy and quantitative analysis of a living biofilm under local irradiation at a defined light dose. In addition, we developed a theoretical model for light-biofilm interaction that accounts for the threedimensional structure of the bacterial biofilm. To inform our model, we examined the light doseresponse in isolated cells and found a profile indicative of a multi-target mechanism of lethality. By comparing the experimental and theoretical results, we identified a loss in killing efficiency as the biofilm grows, due in part to the increase in thickness of the living material inherent to this mode of development. Our findings also highlight a reduction in the intrinsic bacterial sensitivity to blue light as biofilm development progresses, which we attribute to the low oxygen levels typical of densely populated bacterial environments. These findings reveal new features of the photokilling mechanism and redefine the approach to designing effective antimicrobial photoinactivation strategies by integrating the key physical characteristics of bacterial biofilms.

**Significance Statement:** Awareness of the bacterial world’s global importance is steadily growing in both science and society. Among the critical challenges, the continuing increase in multidrug resistance to antibiotics represents a major public health concern reinforcing the urgency of alternative antimicrobial therapies with photoinactivation as a promising approach. However, its full potential can only be achieved through a better understanding of the involved mechanisms in relevant environments. In this study, we combined experimental and theoretical approaches to investigate the photoinactivation of bacteria within a developing biofilm, the dominant bacterial lifestyle. Our comprehensive analysis sheds light on the mechanisms and limitations of photoinactivation in the fight against microbes, which is essential for designing novel antibacterial phototherapies.

## Introduction

The rapid expansion of antimicrobial resistance is now recognized as a serious public health issue (1), and unconventional strategies to fight against microbes are actively being researched. Recently, antimicrobial blue light (aBL) arose as a potential alternative therapy (2, 3). This approach relies on the presence in microbes of natural photosensitizers such as porphyrins, which are excited to the triplet state upon absorption of blue light in the range of 380 — 420 nm. This generates reactive oxygen species that deactivate locally, damaging a wide range of macromolecules and impairing various cellular functions (4–9). Due to the lack of a single biological target, it was postulated that the emergence of bacterial resistance should be less probable (10, 11). To date, blue light-induced killing has been demonstrated in a wide range of microorganisms (both bacteria and yeast), and several studies have reported that irradiation with light doses in the range of 10 to 100 J/cm^2^, using wavelengths between 400 and 470 nm, decrease survival by several orders of magnitude (5, 7, 12–15). Despite a large variability in results and protocols across different studies (3, 10, 16–18), findings from the last decade generally substantiate the view that blue light reduces bacterial cell proliferation. However, the essential factors contributing to the achieved effects remain unclear, as shown in a systematic analysis of the literature for *Escherichia coli* (19).

In this context, we focused our investigations on biofilms, the prevailing lifestyle of most pathogens (20). These multicellular three-dimensional structures typically adhere to surfaces where cells multiply and are confined in a protective, self-secreted extracellular matrix. Within biofilms, cells encounter physical and chemical conditions that differ significantly from those experienced in their planktonic state. This induces the emergence of specific properties, notably multi-factorial resistance to biocides. For now, the question of which physiochemical factors drive the biofilm response to blue light, and how this is accomplished, is largely unanswered. Despite previous reports of blue light inactivation of pathogenic microorganisms in biofilms (17, 21–26), the absence of standardized illumination configurations, biofilm development specifications, and methodological details for assessing photokilling efficiency have prevented drawing any precise conclusions regarding the key parameters that influence the response of bacterial biofilms to blue light. For instance, a popular way to grow biofilms consists of seeding multi-well plates and allowing the cells to form the biofilm either directly on the walls of the wells, or on pegs immersed in the wells and attached to a removable lid. While these settings facilitate performing numerous experimental conditions in parallel, they promote the formation of a biofilm topography that leads to substantial heterogeneity in light exposure, making the analysis of the results challenging. Additionally, uncontrolled variations in cellular environmental properties arise when cells are grown in disparate settings. Biofilm architecture is another factor that may impact light delivery, which could affect the design of an antimicrobial blue light strategy. These obstacles also hinder accurate comparisons of the sensitivity of planktonic cells and biofilm-dwelling cells.

Within this framework, we focused our interest on the *Pseudomonas aeruginosa* PAO1 model strain, an opportunistic pathogen known to colonize the lungs of cystic fibrosis patients, which has the extended potential to develop antibiotic resistance (27, 28). Our aim was to devise an experiment that could decipher the main factors involved in biofilm response to blue light irradiation, by controlling the biological development of the model, the environment, and light delivery. We therefore opted for a device capable of real-time monitoring of a biofilm grown under flow in a millifluidic channel on a confocal microscope stage. This setup offers complete control over the physicochemical parameters of biofilm development such as applied shear stress, nutrient supply and temperature, and allows for the precise delivery of defined light doses at various stages of biofilm formation within a clear, reproducible and coherent geometry.

Thanks to this device, we can investigate how blue light impacts the overall development of the bacterial community, and how killing efficiency evolves with biofilm expansion. Specifically, we demonstrate here that the antimicrobial efficacy of blue light decreases as biofilm development progresses, resulting in a rapid recovery with negligible impact in the long term. We quantitatively analyzed the killing efficiency and kinetics of aBL-induced cell death to identify the underlying mechanisms, by comparing biofilm response to single isolated cells. To explain these observations, especially the decline in blue light efficiency as the biofilm develops, we formulated a physical model that predicts the dose-dependent bacterial photokilling efficacy by taking into account light absorption in the growing material. Importantly, both experimental evidence and theory consistently point to a decline in aBL efficiency as the biofilm ages. We conclude that thickening of the biofilm throughout its development significantly contributes to the light screening effect. Comparing the model with the experimental data also revealed a decrease in the cell sensitivity constant, which could be related to the intense metabolic consumption of oxygen in these cell-concentrated organizations.

Our results reshape the framework for designing a practical antimicrobial photoinactivation strategy, by incorporating the key characteristics of bacterial biofilms, the dominant bacterial lifestyle whose structure and functions impose constraints on photoinactivation. This must therefore be taken into account in any antibacterial phototherapeutic design.

## Results

### PAO1 biofilm displays a deterministic developmental pattern under flow

The formation of PAO1 biofilm in the millifluidic channel was monitored by imaging the internal top surface of the channel where it preferentially forms (Figure 1a); growth kinetics were then drawn from time-lapse imaging of the biofilm in transmitted light. The *µOD* signal (Figure 1b) shows that PAO1 biofilm developed deterministically according to three distinct kinetic regimes, as indicated by the time derivative (Figure 1c). The first phase, lasting approximately 5 hours, follows an exponential growth with an apparent growth rate 1.1 h obtained from adjusting the data to

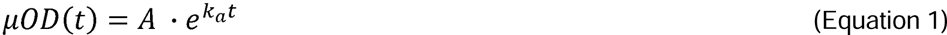

**Figure 1.**
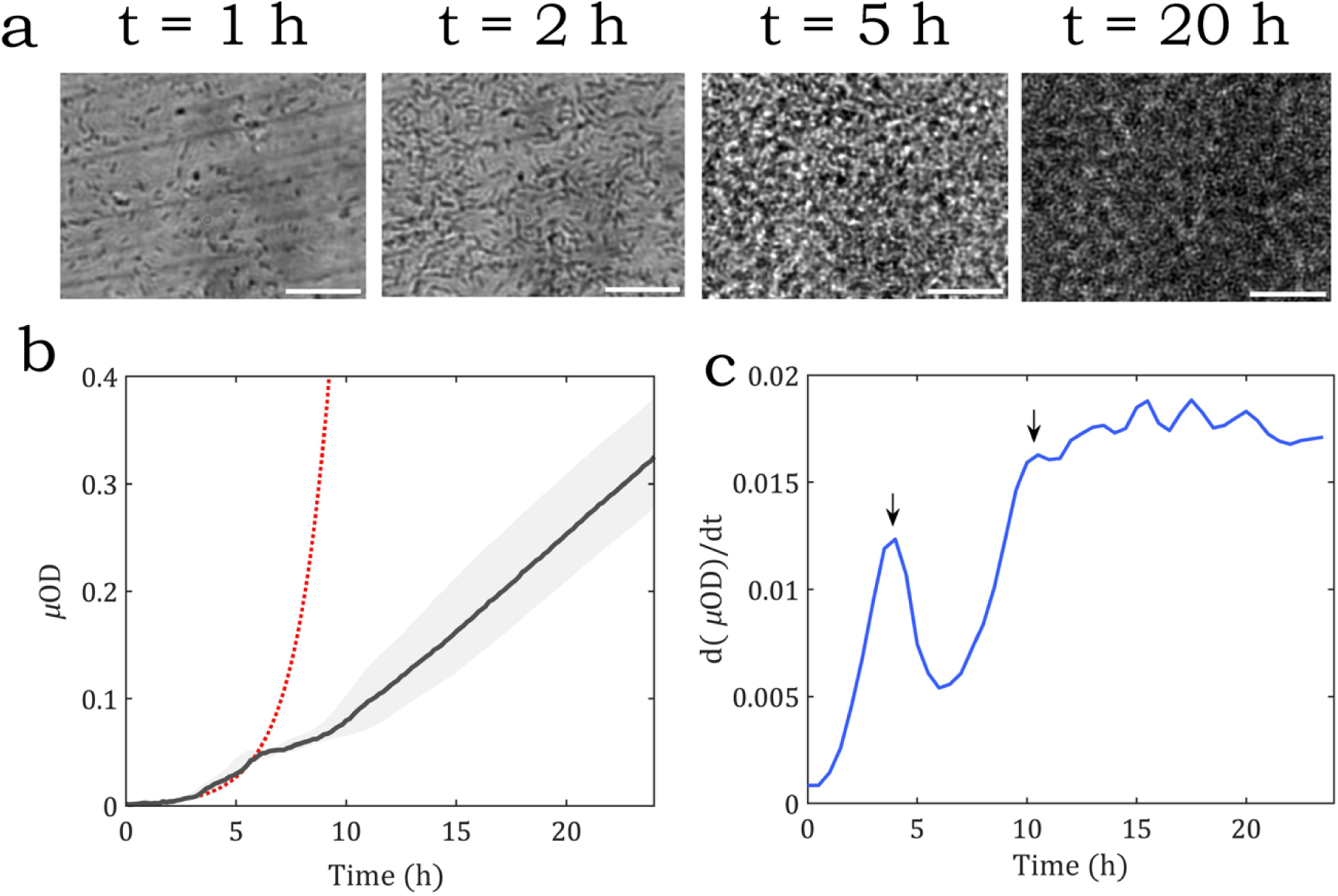
PAO1 biofilm develops under flow according to three distinct phases. (a) Representative brightfield images at characteristic biofilm growth times (0, 2, 5 and 20 hours). (b) Biofilm growth reported by µOD kinetics taken from time-lapse movies. The experimental curve in black represents the average of five distinct positions in two different channels; the standard deviation is shown in the grey shaded region. The initial part of the curve from t = 0 to 6 h has been adjusted to an exponential growth, represented as the red dotted line. (c) The derivative of µOD = f(t) highlights the three distinct biofilm phases, delineated by the maxima marked by the black arrows at t = 5 h and t = 10 hours.

where *A* is the initial *µOD* at *t* = 0 related to the initial number of cells. The biofilm growth then experienced a transition consisting of a transient slowdown followed by a new increase up to a stable value that was reached after approximately 10 h, marking the beginning of the third phase, characterized by a constant rate. Interestingly, microscopy images show that the first kinetic switch correlates with the completion of the first cell surface monolayer (Figure 1a, Movie S1 and S2). After this time point, the cells start to build additional layers and the biofilm thickens. This quantitative description of biofilm development facilitated investigating the biofilm response to blue light in the different developmental stages, from the initial exponential growth to the linear temporal development of the community.

### Blue light killing efficiency declines throughout PAO1 biofilm development

In order to evaluate the aBL killing efficiency on the growing PAO1 biofilm, we irradiated defined positions at different times following the initiation of the biofilm, using a 2007 J/cm^2^ light dose.

Time-lapse imaging of the irradiated zone was recorded in parallel with control non-irradiated positions in the same channel (Figure 2a). The resulting growth curves in Figure 2b show that irradiating the young biofilm in its initial exponential growth phase (within the first five hours) instantaneously hinders the biofilm growth in the captured area, inducing a growth delay of several hours (see Figure 2c).

**Figure 2:**
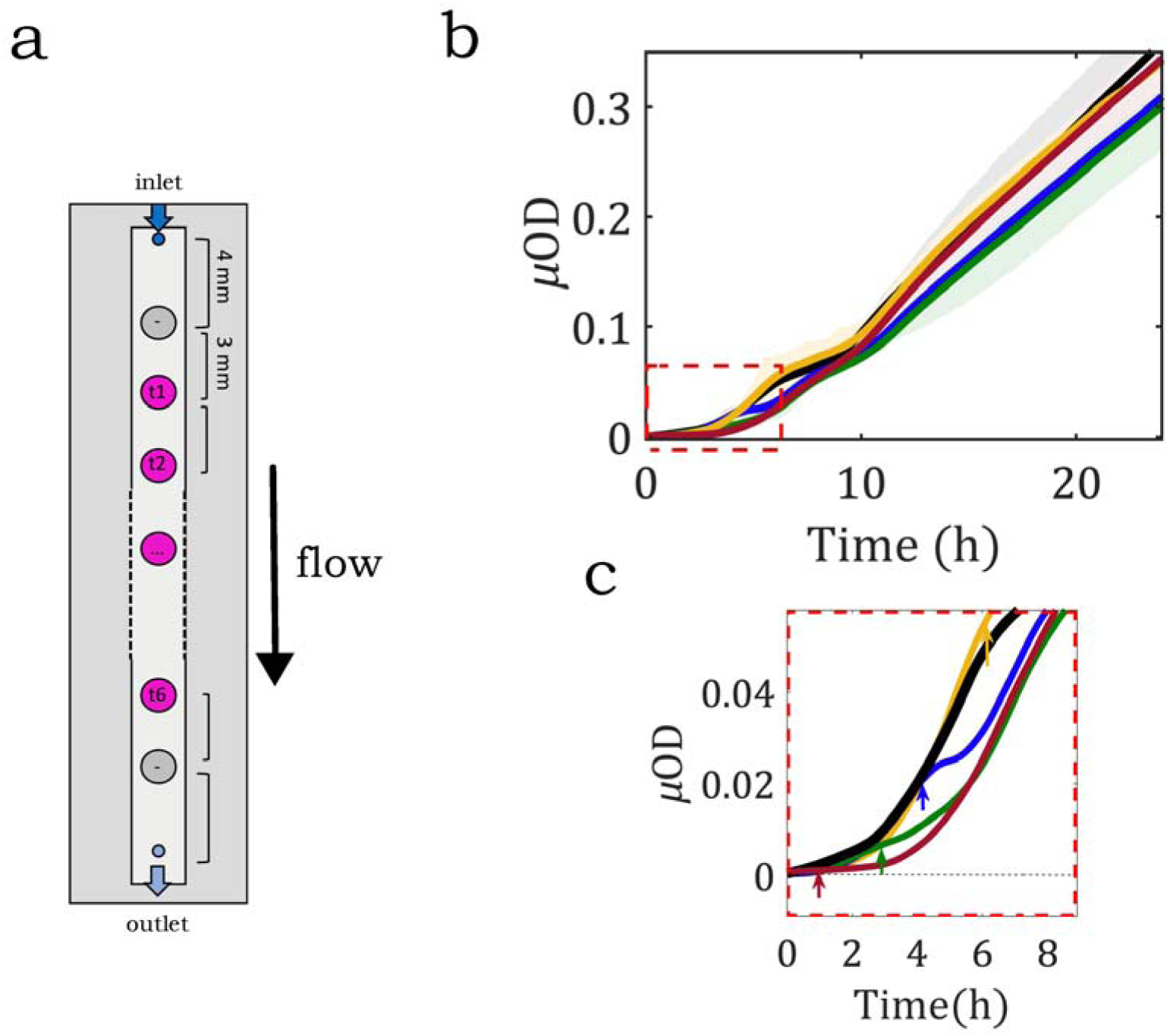
Local irradiation of a PAO1 biofilm affects young biofilm growth. (a) Channel sketch where six of the eight positions were illuminated at 405 nm. (b) Local growth curves; with the exception of the controls (in black), all positions received the same dose (2007 J/cm2) at different times of biofilm growth. The initial time (t = 0) corresponds to nutrient flow start (0.5 mL/h). Irradiation at 1 h is shown in red, at 3 h in green, at 4 h in blue, and at 6 h in yellow. Panel (c) (marked by a red dotted line) focuses on the initial exponential growth phase, with arrows indicating the light dose delivery time of each curve. The curve of each time is the average of at least two independent positions, and is shown shaded with the standard deviation of the data set.

The irradiated zones subsequently recover and essentially match the biomass and growth rate of the controls, demonstrating a limited long-term effect on global biofilm development. To better assess the damage induced by the different irradiations, we measured the local cell mortality through the use of propidium iodide (PI), the cell death marker, in the nutrient flow and collected time-lapse transmission and red fluorescence images throughout biofilm growth, both before and after illumination (Figure 3a). Upon irradiation, PI fluorescence intensity (*F_PI_*) suddenly increased up to a stable plateau, according to a first order kinetic (Figure 3b and Movie S3 and S4) as follows:

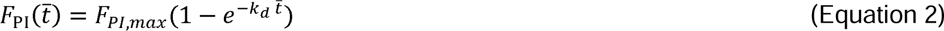

**Figure 3:**
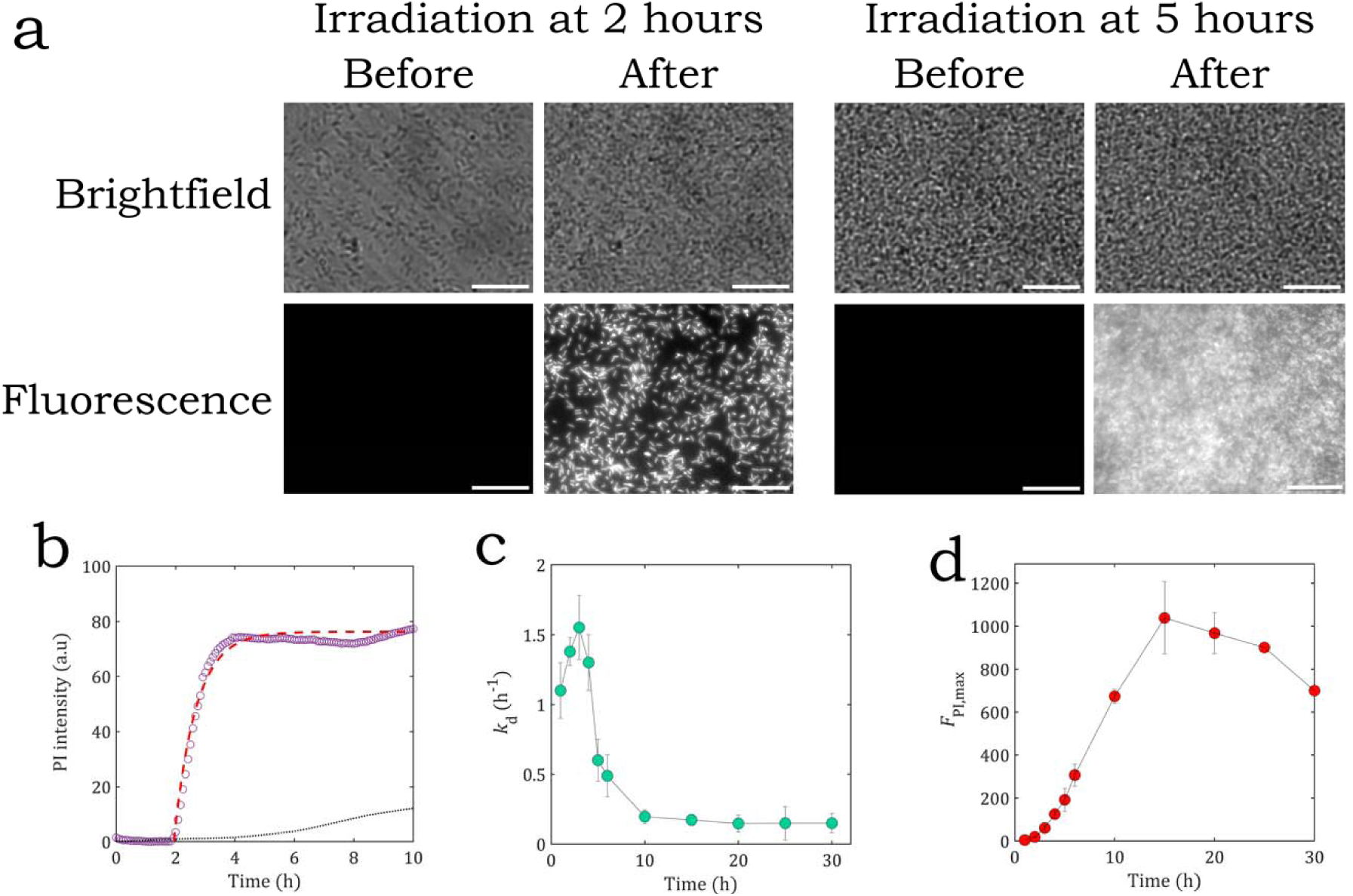
Blue light toxicity during biofilm development. (a) Transmission (upper row) and red fluorescence (lower row) microscopy images were recorded just before (left column) and after (right column) irradiation (2007J/cm2) of the growing biofilm supplied with PI (3 µM). Fluorescence images were taken at the PI fluorescence plateau; the white scale bar represents 25 µm. (b) Measurements of PI rise kinetics (purple dots) were adjusted to the first order equation (Equation 2; red dashed line). The evolution of kd (c), the kinetic constant for PI increase in the biofilm, and (d) FPI,max as a function of the time of irradiation.

where *F_PI,max_* is the maximum intensity at the kinetic plateau, related to the number of dead cells *N_d_*; is the time elapsed from irradiation triggering; and *k_d_* is the kinetic constant that provides the characteristic time for the emergence of PI-labelled cells. The PI-DNA interaction occurs in less than two minutes in pre-permeabilized cells, indicating that here the full cell permeabilization rate-limiting step, in which the detrimental effects of blue light result in cell death, has been identified. *F_PI,max_* and *k_d_* were obtained by fitting time-lapse data to Equation 2, and are plotted as a function of biofilm age at the time of irradiation (Figure 3c-d). *k_d_* drastically decreases for irradiations performed after 5 hours of biofilm development, revealing that light-induced killing takes more time to complete as the biofilm matures beyond the initial phase. *F_PI,max_*, which reflects the absolute number of dead cells, displays a non-monotonic profile that can be understood only relative to the total biomass at the irradiation time.

We therefore defined the killing efficiency *K_E_* as *K_E_ = N_d_/N_0_*, where *N_d_* is the number of dead cells and *N_0_* is the total number of cells at the start of the irradiation given by the *µOD* signal calibrated by GFP fluorescence at a short time scale (see Materials and Methods). The results in Figure 4 indicate that *K_E_* remains stable and near a value of 1 for irradiations applied within the first five hours of development and then declines gradually and consistently, similar to the trend observed for *k_d_*. Interestingly, the shift in the curve at about 5 h coincides with the end of the biofilm’s initial growth phase and the point at which the community begins to expand three-dimensionally after completing the first monolayer. These results led us to hypothesize that the 3D structure of the bacterial biofilm and its thickening during growth could be responsible for the observed reduction in blue light efficiency. To test this idea, we devised a simple mathematical model to describe the light screening effect and the potential light dose reduction induced by the addition of new cell layers as the biofilm grows.

**Figure 4:**
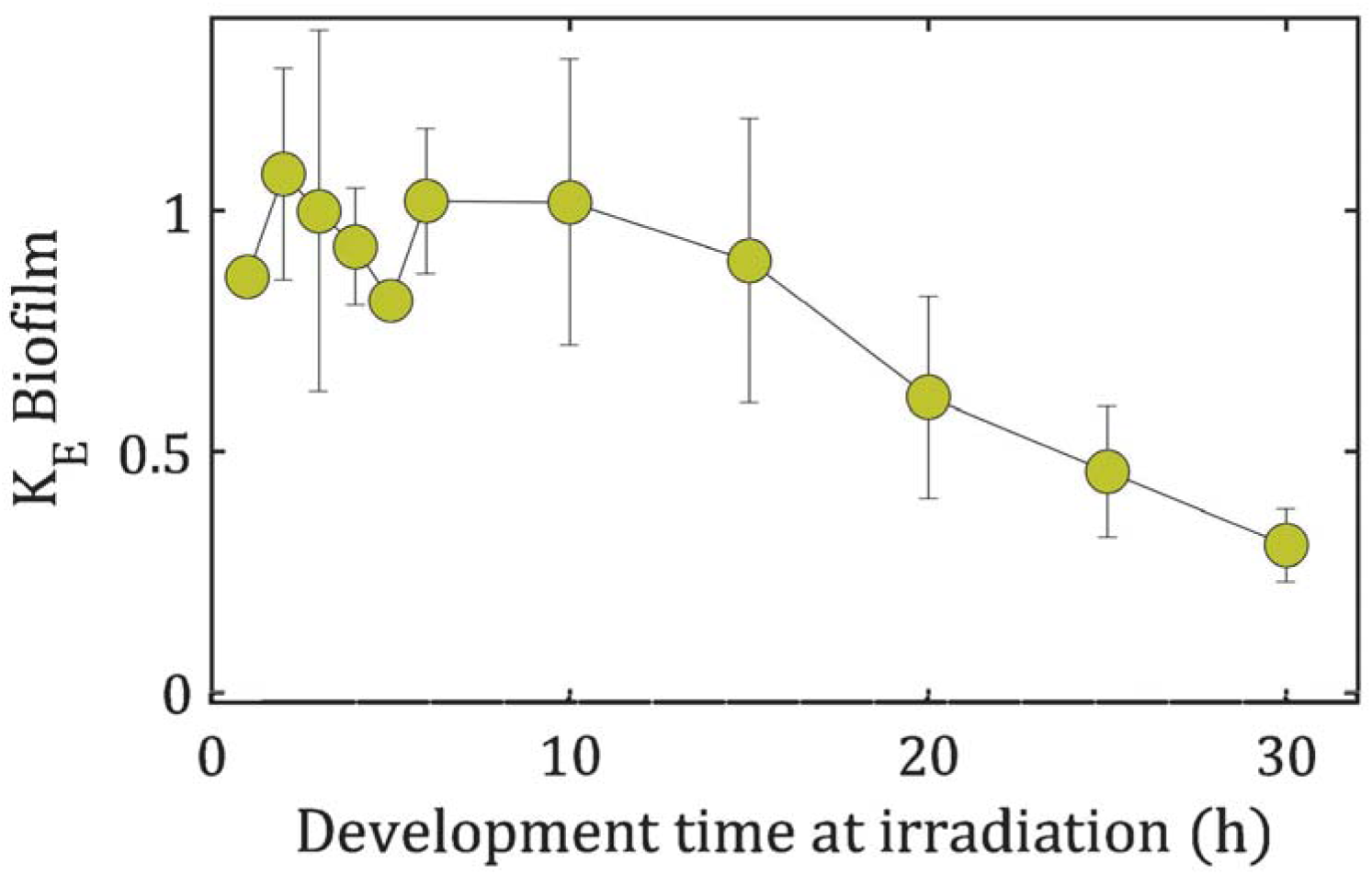
Killing efficiency decreases with biofilm aging. To examine killing efficiency, KE = Nd/N0 was derived from FPI,max and biofilm µOD for irradiations at increasing biofilm development times. Irradiations were performed at 2007 J/cm2 under constant nutrient flow (0.5 mL/h) and in the presence of PI (3 µM).

### Modeling light dose attenuation induced by biofilm thickening

To evaluate the light dose attenuation caused by biofilm thickening, we considered the light absorption process as described by the Beer-Lambert law, which states that the transmitted light intensity impinging orthogonally on the biofilm decreases exponentially with the optical path length and concentration of absorbing compounds, yielding:

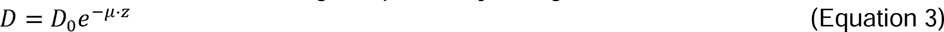

where *D* is the light dose corresponding to the energy fluence impinging on the unit surface orthogonal to the illumination direction at a given depth *z* inside the biofilm; *D_0_* is the light dose at *z* = 0; and μ represents the effective absorption coefficient, an intrinsic invariant of the cell. This coefficient is related to the presence of various absorbers composed of cellular material, including the membrane and the intracellular medium packed with proteins and DNA, and it can be expressed by the formula ∑, in which and respectively represent the molar extinction coefficient and the concentration of each individual absorber. To facilitate comparisons between the model and the experimental data, we chose to express *z* in terms of the number of cell layers, with μ representing the effective absorption coefficient per unit cell layer. To evaluate this, we measured light transmission through small patches of PAO1 cell monolayers confined in an agar pad illuminated by a 405 nm LED and obtained a value of μ = (0.22 ± 0.1) per cell layer (Supplementary Information I and Figure S1). In order to relate this to the killing efficiency in the biofilm, we must also presume a theoretical light dose dependence of the blue light intrinsic killing efficiency for a population of isolated bacteria.

### Modeling light dose-dependence of aBL intrinsic killing efficiency

Despite the large number of reports dedicated to aBL (1–19, 21–26), studies on the light dose-dependence of the killing efficiency are sparse and poorly generalizable (15). In order to infer the dependence of the killing efficiency on the light dose, we devised an experiment aimed at specifically measuring the dose-response of a population of isolated cells that do not form a biofilm under our defined conditions. For this, we employed PA01-GFP cells deposited in agar pads with added PI, which allows monitoring light irradiation toxicity at the different applied light doses. This configuration places all of the cells in a unique optical plane, which enables defining a single light dose value for all of the cells and precisely monitoring cell death. The photokilling efficiency as a function of the delivered light dose is reported in Figure 5. The curve shows a pronounced inflection at low light doses, suggesting the presence of a threshold effect.

**Figure 5:**
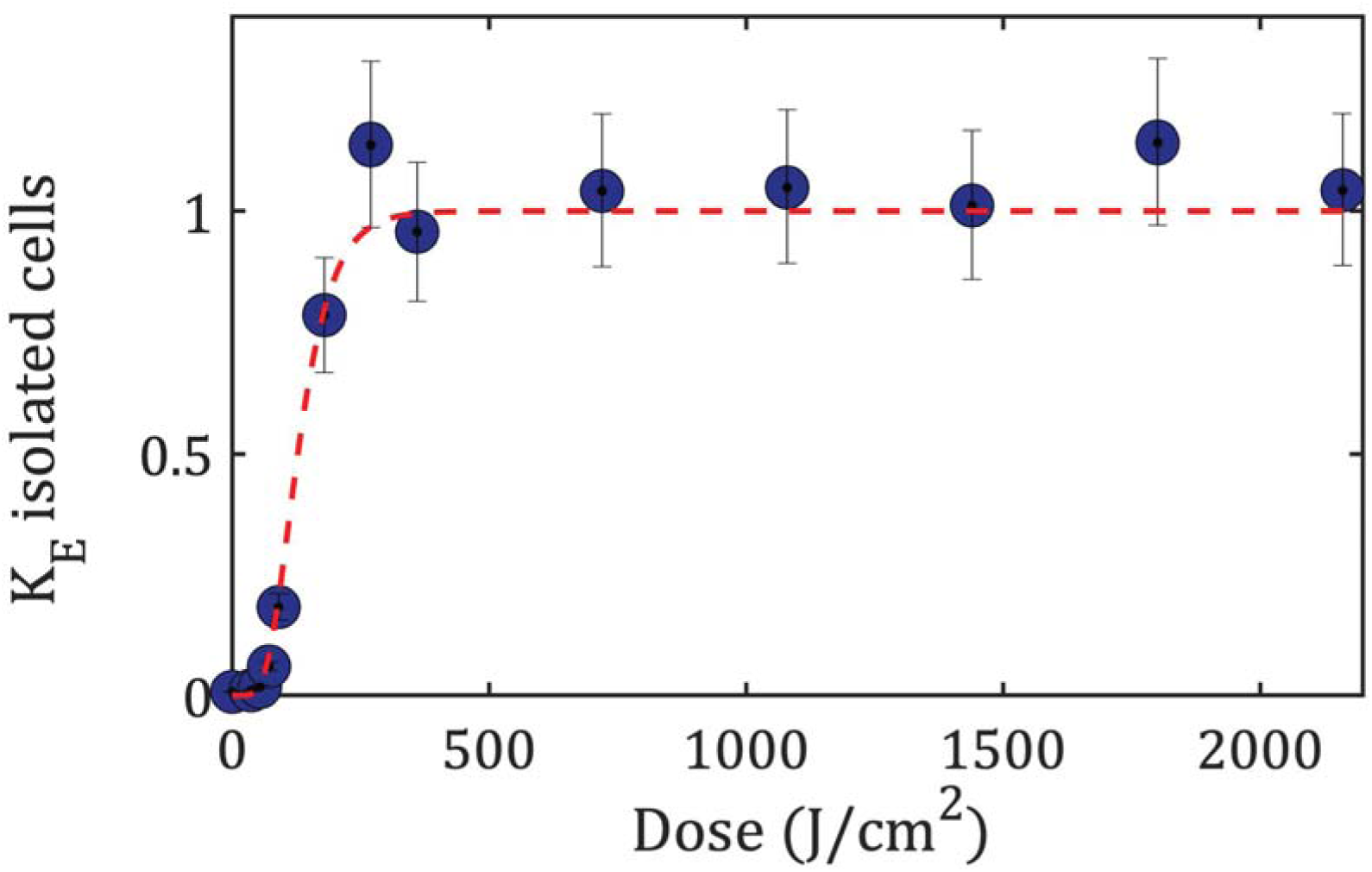
Isolated cells display an S-shaped sensitivity to blue light. Killing efficiency is illustrated as a function of light dose (405 nm) in isolated cells immobilized in an agar pad. Experimental data (blue dots) are shown with their adjustment to Equation 5 (1) (dashed red line), providing ks = (0.021 ± 0.003) (J/cm2)-1 and n = (10 ± 2).

To account for this result, we presumed a dose-dependence similar to what has been proposed for modeling ionizing radiation activity on living cells, assuming a multi-target mechanism (29). This model considers radiation as a sequence of projectiles — here, impinging photons — that produce hits with a Poisson law probability, and assumes that a single hit damages a single target per cell with a probability P_1_. The single-target damage is assumed to be sublethal with a probability equal to

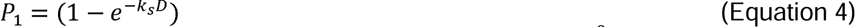

where *D* is the energy per unit surface expressed in J/cm^2^ that is related to the number of photons per unit surface during the whole illumination time, and *k_s_* is a sensitivity constant that is independent of *D*. Thus, assuming that cell death is induced by hitting *n* targets per cell, its probability is *P_n_* = (*P_1_*)*^n^*. The killing efficiency is therefore:

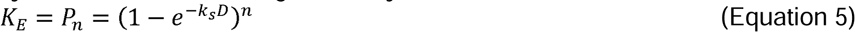

By fitting the isolated cell killing efficiency data (Figure 5) with Eq. 5, the best agreement between the data and the model is obtained for *k_s_* = (0.021 ± 0.003) (J/cm^2^)^-1^ and *n* = (10 ± 2).

Notably, the multi-target model is compatible with the living cell population surviving a dose *D* that exponentially decreases with *D* (29), in which the *k_s_* and *n* parameters are independent of *D*.

### Modeling aBL killing efficiency in a thickening biofilm

To take into account the exponential decrease in light dose with the optical path length and the consequent decrease in photokilling efficacy, we generalized the expression of the killing efficiency established for isolated cells (Eq. 5) to a growing biofilm of thickness *H* to obtain the following formula:

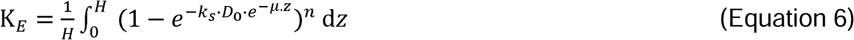

in which killed cells have been integrated over *H*, assuming constant cell volume density. Numerical integration of this equation was used to simulate the model and examine the dependence of the killing efficiency on the key system parameters *k_s_*, *D_0,_ n*, and μ (Figure 6a, b, c and d).

**Figure 6:**
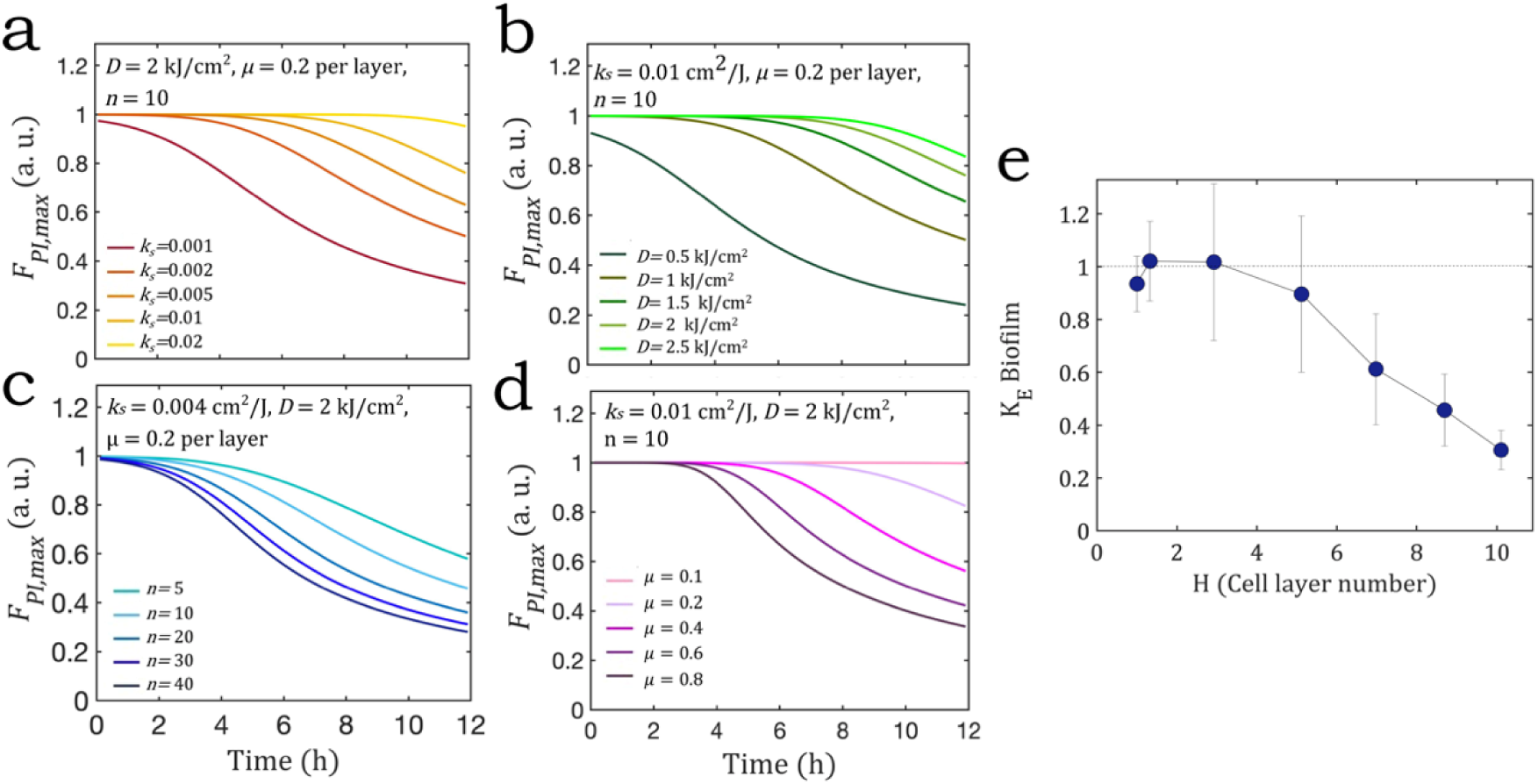
Killing efficiency calculated from the model for different sets of parameter values. (a) 0.001 < ks < 0.02 (J/cm2)-1 with D = 2 kJ/cm2, µ = 0.2 per layer, and n = 10. (b) 0.5 < D < 2.5 kJ/cm2 with ks = 0.01 (J/cm2)-1, µ = 0.2 per layer, and n = 10. (c) 5 < n < 40 with D = 2 kJ/cm2, µ = 0.2 per layer, and ks = 0.01 (J/cm2)-1. (d) 0.1 < µ < 0.8 with D = 2 kJ/cm2, ks = 0.01 (J/cm2)-1, and n = 10. (e) The experimental killing efficiency is shown as a function of the number of cell layers, obtained using a light dose of 2007 J/cm2.

Figure 6 shows the behavior of the model according to a series of parameters selected for their experimental significance, in the range of one to about ten cell layers. The trend is qualitatively consistent with experimental observations showing that killing efficiency tends to decrease as the biofilm ages, and thus as it thickens (Figure 6e). These simulations illustrate the different parameters that can affect the killing efficiency. Logically, the loss of killing efficiency intensifies with increasing extinction coefficients, but declines with stronger doses and higher sensitivities. Additionally, a greater number of targets needed to induce lethality are associated with an amplified loss of efficiency during biofilm development.

To quantitatively compare the model with the data, we converted the experimental data giving the killing efficiency as a function of the biofilm development time, i.e. *K_E_* = *f(t)*, into data giving the variation of the killing efficiency with *H*, i.e. *K_E_* = *f(H)* (Figure 6e), in which *H* denotes biofilm thickness expressed in number of cell layers. The conversion law used to change the variable *t* into variable *H* is provided by the biofilm *µOD*, assuming that the absorbance increases in proportion to the number of layers (Supplementary Information II and Figure S2). We therefore quantified the agreement of the theoretical killing efficiency function (Equation 6) with the experimental data using the Chi-squared (^2^) test as an estimator of goodness-of-fit (see Supplementary Information III and Figure S3 for a detailed explanation of the numerical procedure).

We constrained the minimization by setting *D_0_* to an experimentally determined value (*D_0_* = 2007 J/cm^2^) and varied *n* and *k_s_* for a series of *µ* values between 0.2 and 0.3 per cell layer, as indicated by the experimental determination (*µ* = 0.22 per layer). We thus observed a well-defined, crescent-shaped area in which the ^2^ value is minimal and outlines a parameter set domain consistent with the experimental results (Figure 7a). A minimal ^2^ value of 1.75 was obtained for *µ* = 0.26 per cell layer, which returned a pair of *k_s_* and *n* values equal to 0.004 (J/cm^2^)^-1^ and 14, respectively. Using this parameter set, we compared the time dependence of the killing efficiency derived from the model with that given by the experimental data (Figure 7b) and obtained a good agreement within the limits of the error associated with the experimental measurements.

**Figure 7:**
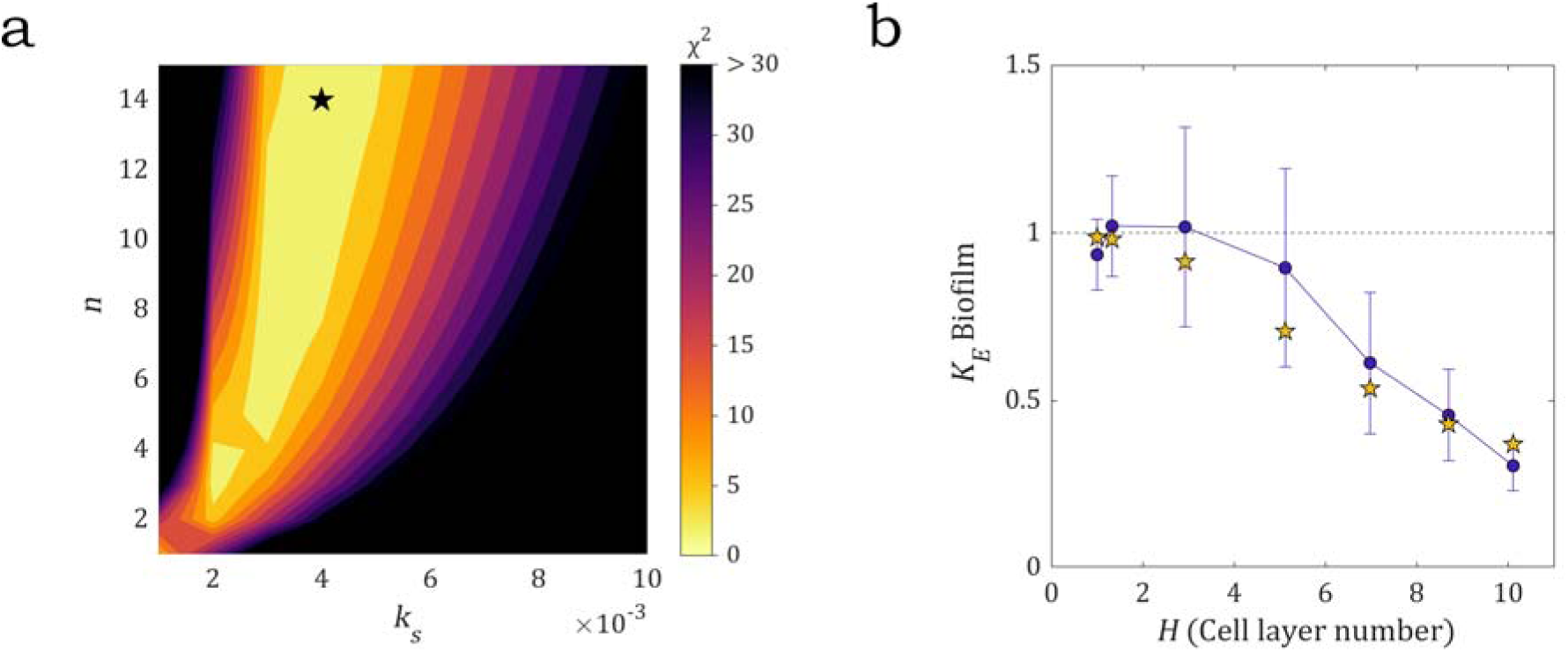
2 contour map. (a) 2 values reported for the n and ks parameters ranging between 1—10 and 1—10 10-3 (J/cm2)-1, respectively calculated for µ = 0.26 per cell layer and a light dose of 2007 J/cm². The black star indicates the parameter set corresponding to the minimum 2 value, i.e. ks = 0.004 (J/cm2)-1 and n = 14. (b) The experimental data with associated errors (blue markers) vs. the theoretical model (yellow markers), calculated with parameters corresponding to the black star in (a).

Interestingly, parameters *µ* and *n*, which were filtered out by ² minimization, are in reasonable alignment with the experimental determination. Our experimental value of *µ* = 0.22 per cell layer is sufficiently comparable to the value of 0.26 returned by the model. Moreover, the *n* value of 10 obtained in the isolated cells experiment is on the same order of magnitude as the value of 14 determined by the ² minimization. By contrast, the model revealed a significantly lower sensitivity constant (by a factor of 5) than the value determined in the isolated cell experiments. This result suggests that, in addition to the light dose attenuation caused by biofilm thickening, a decrease in cell intrinsic sensitivity occurs as the biofilm grows, participating in the loss of efficiency. This is in line with the view that the biofilm lifestyle induces significant environmental changes.

## Discussion

We report here a quantitative analysis of the antibacterial effect of blue light on a *P. aeruginosa* biofilm grown under constant flow of growth medium in a millifluidic device, which enabled characterizing the spatiotemporal development of the community in real time. Our results demonstrate that localized irradiation of the biofilm induces a transient delay in biofilm development due to photokilling, which is eventually outpaced by the growth of neighboring untreated populations. Significantly, this demonstrates that biofilms often exhibit complex topographies, making it unrealistic to irradiate the entire bacterial population.

By looking more closely at the induced phototoxicity, we provide evidence that aBL efficiency decreases throughout biofilm development. In order to clarify the factors behind this decrease, we have proposed a theoretical model based on the cell intrinsic aBL dose-response and material absorption laws.

Irradiating isolated cells in a configuration where the cells receiving a defined dose of photons could be monitored in real time revealed a killing mechanism in which n targets must be damaged to achieve irreversible toxicity. This view, which exquisitely fits the experimental data, is in line with previously established radiation theories that modeled the effect of X-rays on *E. coli* (30). Even if this theory was developed in relation to ionizing radiation, we considered extending its applicability to the optical radiation field, as the model is ultimately based on cell-localized damage resulting from single-photon interaction effects. The model that considers light as quanta of projectiles better describes the experimental behavior than previous modeling dedicated to phototoxicity, which mainly focuses on aBL inactivation kinetics (31, 32). To the best of our knowledge, few studies have addressed the question of light dose-dependence of the killing efficiency by also incorporating the key characteristics of bacterial biofilms such as their threedimensional nature and specific physico-chemistry.

Previous works reported by Kumar and collaborators, have transposed the kinetic models to fit the killing efficiency data, using for instance a modification of the Gompertz growth equation (15). However, these models do not intrinsically extrapolate to zero and the function must be forced to the origin by subtracting a constant that has no physical meaning. Therefore, although a satisfying adjustment of the experimental data can be obtained, the modeling does not support any mechanism of action or predictive viewpoint. The model we propose here indicates a minimum lethal number of targets per cell, on the order of 10 for isolated cells.

In our experiments, we determined a cutoff dose on the order of 50 J/cm^2^, which corresponds to a photon flux of approximately 10^10^ photons/s per cell provided during about 100 s. This indicates an extremely unfavorable ratio of photons/killed cell, which likely arises from photosensitizer-limited availability, oxygen-limited concentration, antioxidant processes, and ROS production quantum yield. In fact, it can be calculated from a previous study (6) that about 160 molecules of photosensitizer are expected per cell in *P. aeruginosa*. It is also known that bacteria possess antioxidant defenses that may reduce the light damaging efficacy by deactivating oxygen radicals (3, 6). These numbers should be considered in relation to the mean intensity of the solar illumination at the Earth’s surface, e.g. 140 mW/cm^2^ according to previous work (33). This indicates that the lethality threshold could be reached within approximately 6 minutes of sunlight exposure, suggesting that dark habitats as well as the biofilm life mode should be more favorable to these bacteria.

We determined that light attenuation due to the thickening of the biofilm during its growth partially explains the loss in efficiency observed as the biofilm ages. The sets of parameters returned by fitting the experimental data with the model are compatible with the idea that bacterial development under a biofilm lifestyle also induces a decrease in the bacteria photosensitivity constant. This result can be related to the rapid decrease in oxygen levels that occurs in a biofilm, due to the high consumption in a densely populated cellular environment, as previously demonstrated in an *E. coli* biofilm through the evolution of GFP fluorescence (34). This hypothesis is supported by our work, since we also observed in our *P. aeruginosa* biofilm that the collinearity between GFP and biomass begins to diverge as the biofilm enters the multilayer stage. Such a decrease is very likely to negatively impact the aBL killing efficiency by impairing ROS formation, consistent with the *k_s_* decrease revealed by the data modelization.

The model also returned a *µ* value slightly higher than the measured one. This might be due to the fact that we approximated the biofilm to an ideal material composed of purely ordered stacks of absorbing cells, whereas a real biofilm possesses an extracellular polymer matrix and likely exhibits some disorder in the piling of the different layers. However, the significantly lower molecular density of the extracellular matrix compared to the cells justifies neglecting its contribution to the overall biofilm absorption. We also estimated that structural disorder was not expected to significantly change our estimated absorption. Indeed, previous numerical work (35) shows that switching the order parameter value from 0 to 1 causes a reflectance increase of about 20%, which might explain the observed discrepancy.

We conclude that two simultaneously occurring hurdles affect bacterial photokilling in biofilms as they grow: the increasing thickness of biological material, which screens the impinging photons; and the potential onset of hypoxia, which hinders ROS formation — the primary mechanism of photokilling in bacteria. In this context, effective phototherapy strategies should focus on targeting young monolayer biofilms to avoid any screening effects, and also to mitigate hypoxia by ensuring direct access to environmental O_2_, so as to balance consumption. Nonetheless, even suboptimal light-induced killing of a mature biofilm could be valuable for future clinical applications, especially when combined with antibiotics. Moreover, collecting information about bacterial target-specific optical properties will undoubtedly help in designing more efficient phototherapy protocols.

Our approach can easily be extended to any irradiation wavelength other than blue, and also to photokilling in the presence of an external photosensitizer, provided that certain changes are made (e.g. measuring the biofilm absorption that corresponds to the irradiation wavelength, etc.). This will greatly expand the applicability of both the experimental methods and the theoretical model. Finally, our study demonstrates the depth and effectiveness of combining experimental and theoretical approaches to generate new insights and practical strategies for antibacterial phototherapies.

## Materials and Methods

### Chemical reagents, bacterial strains and culture media

Propidium iodide (Sigma-Aldrich) stock solutions were prepared from powder in distilled water for a stock concentration of 3 mM. M9 medium (Na_2_HPO_4_ 12.8 g/L; KH_2_PO_4_ 3 g/L; NaCl 0.5 g/L; NH_4_Cl 1 g/L; MgSO_4_ 2 mM; CaCl_2_ 0.1 mM; glucose 4 g/L) was supplemented with casamino acids (2 g/L) to prepare M9CA medium, which was used in all experiments.

*Pseudomonas aeruginosa* (PA01) and its green fluorescent variant (PA01-GFP) were kindly provided by the University of Liverpool. PA01-GFP constitutively expressed GFP, as previously described (36). Overnight cultures were obtained by inoculating a smear from the frozen stock in 5 mL of M9CA medium and incubating at 37°C, under constant agitation (500 rpm). For biofilm experiments, the overnight culture was diluted in 5 mL of fresh M9CA medium to an optical density of 0.1, then incubated for 2 hours under the same conditions (37°C, 500 rpm). This exponential culture was then diluted to an optical density of 0.1, corresponding to ∼10^7^ CFU/mL.

### Agar pad preparation

A 1.5% low-gelling agarose solution in M9CA medium, supplemented with propidium iodide (PI) at a final concentration of 3 µM when needed, was added to a spacer-delimited cavity fixed to a glass slide (125 µL frame, 1.7 cm by 2.8 cm; Thermo Fisher Scientific), which was smoothed by carefully sliding a coverslip on the free surface before depositing bacterial cells (2 µL of an exponentially growing suspension with an OD of about 0.1) and closing the device by sticking a coverslip on the spacer.

### Millifluidic device

Millifluidic channels molded in PDMS were microfabricated as previously detailed (37). Growth medium was continuously supplied using syringe pumps. For connections, we used stainless steel connectors (0.013” ID and 0.025” OD) and microbore Tygon tubing (0.020” ID and 0.06” OD) supplied by Phymep (France). Channel dimensions were 30 mm x 1 mm x 0.25 mm, providing an inner volume of 7.5 µL.

### Biofilm formation

An exponentially growing culture diluted in M9CA medium at an OD of 0.1 was injected by syringe directly into the PDMS channels before connecting the tubing. The cells were then incubated inside the channel for 30 minutes at 37°C. M9CA was then continuously pushed into the channels at a rate of 0.5 mL/h throughout the entire duration of the experiment. The start of the flow defined time zero (t = 0) for all experiments.

### Microscopy imaging

We used an inverted NIKON TE300 microscope equipped with a motorized x, y, and z stage. Images were collected using different objectives: Nikon 4X Plan (NA 0.10 WD 30 mm), 40X S Fluor (NA 0.90 WD 0.11-0.23 mm), and 40X Plan Fluo (NA 0.90 WD 0.11-0.23 mm). Brightfield images were collected in direct illumination (no phase). Fluorescence acquisitions were performed using either the green channel filters for GFP (Ex. 482/35, DM 506 Em. FF01-536/40) or the red channel filter for propidium iodide (PI) (Ex 562/40nm DM 593 Em 641/75). Excitation was performed using LED lines (CoolLed pE-4000) at 490 nm or 555 nm (50% power level), and exposure times of 500 ms or 100 ms for the green and red emissions, respectively. A Hamamatsu ORCA-R2 EMCCD camera was used for time-lapse acquisitions of 1344 x 1024 pixel images to capture an xy field of view of 215 µm x 165 µm. Brightfield and fluorescence images were typically collected for 24 hours at a frequency of 6 frames per hour.

### Irradiation and light fluence measurements

*In situ* irradiations of the biofilm at different developmental times were performed on the microscope stage using a 405 nm laser diode (LDI-7, 89 North, VT, USA) and a Nikon Plan Fluor 40x/0.60 WD 3.7-2.7 mm objective. The light power was measured with a photodiode (S130VC, ThorLabs) connected to a digital power meter (PM100D, ThorLabs) at the objective focal plane level. The biofilm imaged area received a mean irradiance of ∼18 W/cm^2^, measured at 100% of the laser capacity and according to the method detailed in Supplementary Information IV and Figure S4. The dose was adjusted by varying the irradiation time.

### Microscopy quantitative descriptors

Micro-optical density (*µOD*) was measured from transmitted light images according to *µOD* = ln (*I_0_/I*), by analogy with the Beer-Lambert law where *I_0_* and *I* indicate the incident and transmitted light, respectively. *I_0_* and *I* are given here by the intensity per pixel averaged over the entire image recorded in a channel containing only medium or the growing biofilm, respectively. As previously reported, this quantity proportionally reports the bacterial biomass developing in the channel as long as no signal saturation occurs, i.e. *µOD* < 0.7 (37).

The GFP fluorescence signal (*F_GFP_*) was recorded in parallel with *µOD* on PAO1-GFP biofilms. In the early stage of biofilm development, constitutive expression of the protein provides a fluorescence signal proportional to the number of cells *N* (34). The GFP images acquired during the first 3 to 4 hours of the experiments were used to delineate single cells and determine their count, *N* (number of cells per image). Then, we calculated the image mean fluorescence per pixel fluorescence, *F_GFP_*, by subtracting the background *b_g_*, obtained by recording an image in the absence of cells from the raw image intensity *I*, giving *F_GFP_* = *I - b_g_*. The fluorescence weight *w_i,_ GFP* was deduced as *w_i, GFP_* = *F_GFP_/N* (see details in Supplementary Information V, Figure S5 to S7). The absolute number of cells, *N*, was used to calibrate the *µOD* signal.

### Killing efficiency evaluation

To evaluate cell death induced by irradiation, the growth medium was supplemented with propidium iodide (PI) at a final concentration of 3 µM continuously since the initial step of the biofilm formation. Only cells with compromised membranes permeable to PI could be labeled and detected using the red fluorescence path (38). We verified that the exposure of cells to PI did not affect the viability or development of the PA01 and PA01-GFP biofilms (Supplementary Information VI and Figure S8).

The PI fluorescence signal (*F_PI_*) was recorded throughout the entire duration of biofilm development using time-lapse imaging to evaluate the number of dead cells. Similar to GFP fluorescence, the single-cell PI unit fluorescence weight *w_i, PI_* was derived from images in which the cell population was low enough to delineate and count individual cells. The number of dead cells *N_d_* in larger biofilm populations can therefore be deduced from PI intensity (*F_PI_* = *I_PI_ - b_r_*), with *I_PI_* representing the raw image intensity and *b_r_* the red background intensity in the absence of dead cells, according to *w_i,PI_* = *F_PI_/N_d_* (Supplementary Figure S4).

## Acknowledgments

This work was supported by the project “Light4Lungs”, H2020-FETOPEN-2018-2020, Grant Agreement n.863102, by the International Emerging Action program of CNRS and by the European Union – Next Generation EU - National Recovery and Resilience Plan, Mission 4 Component 2 - Investment 1.5 - THE - Tuscany Health Ecosystem - ECS00000017 - CUP B83C22003920001. The authors thank the members of the Light4Lungs group for fruitful discussions on this work.

## Data availability

All data will be available on demand We are currently in the process of getting sufficient storage space for our data repository at CNRS Research data center (https://entrepot.recherche.data.gouv.fr/dataverse/cnrs) that will provide us with a unique DOI for our datasets. This will be ready upon publication.

## Supplementary Information

### Supplementary Information I: Single cell absorption coefficient

Cell absorption coefficient,, has been measured by considering transmission microscopy images (Figure S6). PAO1 cells were deposited on a thin agar layer in between a glass slide and a coverslip, then imaged in brightfield (60x, Nikon). Images are collected in ten different fields of view and the intensity per pixel of the small cell patches are analyzed individually (as shown in the yellow box). The background intensity (I_0_) is measured on cell-free regions of 10 x 10 pixels, while the light intensity transmitted through one cell diameter (I) is considered to be equivalent to what is transmitted through a cell monolayer. Three spots of 3 to 4 pixels are measured over a single cell as shown on the right insert. The results are shown in Table S1. The value of is deduced from these measurements from . Using = 1, we obtain per cell layer.

### Supplementary Information II: Conversion of biofilm development time into biofilm thickness

In order to compare the experimental data with the model we transposed the time-dependent curve to a biofilm thickness-dependent curve where is the thickness of the biofilm expressed in number of cell layers. To build time-thickness correspondence, we rely on the Beer-Lambert law which states that optical density grows linearly with light path length, or equivalently that *µ*OD proportionally varies with the number of cell layers (Figure S2).

### Supplementary Information III: ^2^ test, Numerical procedure

The experimental data report the killing efficiency, with associated absolute error, relative to a set of different values. For each of these values, the theoretical killing efficiency was also calculated using the following formula: at a specific light dose, with the other parameters assuming values from any of the possible combinations obtained by varying the parameters within the following ranges: : varying between 1 and 12 (in steps of 1) : varying between 0.1 and 1.0 (in steps of 0.1) varying between 0.002 and 0.030 (in steps of 0.002 up to 0.012 and then in steps of 0.005 starting from 0.015) The Chi-squared () value was obtained by the following formula: for each combination of the parameters. A typical result of this calculation is shown in Figure S8, showing the values obtained for a light dose of 2007 J/cm^2^ and for = 1 and = 10.

### Supplementary Information IV: Irradiance determination

The mean irradiance and dose values corresponding to 405 nm irradiation were estimated by the following methodology. A fluorescent plastic slide (Chroma, ref. 92001) with a uniform fluorophore density was imaged by a 40x, NA = 0.6 objective (Nikon) with ORCA-Fusion BT Digital CMOS camera (Hamamatsu), taking care to avoid image saturation and photobleaching of the fluorophore. The corresponding image was calibrated to obtain the µm/pixel ratio value by imaging a stage micrometer in the same conditions. The numerical matrix corresponding to the spot image (370 x 370 µm, 2072 x 2072 pixels, 16-bit gray levels) was analyzed by the Origin® software and fitted with the following function: obtained by adding a constant background z_0 to a bi-dimensional Gaussian function with amplitude A, centroid coordinate (x_0_, y_0_) and standard deviation w. Following this, we derive the background- free and normalized function (over the whole x-y plane):

To properly quantify the amount of power received by the area imaged in the experiments (region B, area S(B) = 215 µm x 165 µm, Figure S4), we calculate the integral of f(x,y) over the imaged area. Knowing that the illumination spot is centered with respect to the imaged region B (meaning x_0_ = 0 and y_0_ = 0), we evaluate the following integral over B:

This means that the biofilm imaged region receives 42.5% of the whole spot power P. This last was measured by a power meter at the exit of the microscope objective (40x) and corresponds to 15 mW. The mean dose was then calculated by D = 0.425 x P x t_irr_ / S(B), being t_irr_ the irradiation time and S(B) the imaged region area. For t_irr_ =111 s we obtain D = 2007 J/cm^2^.

### Supplementary Information V: Image Analysis

Transmitted light and fluorescence movies provide optical signals as a function of time. In order to move from these raw data to biofilm parameters, we developed an image treatment and analysis procedure enabling to convert the micro-optical density and fluorescence intensities in cell numbers. From previous work (1, 2), we already know that the micro-optical density (µOD) is proportional to the number of cells building the biofilm up to a value of 0.7 after which saturation occurs. Similarly, GFP fluorescence intensity— when *gfp* gene is under the control of a constitutive promoter as it is the case here — also reports the number of cells at the lowest cell densities at which the consumption of oxygen is not too high. In the linear regime, GFP and µOD signals are colinear.

*GFP images analysis for µOD calibration*.

To calibrate the GFP signals in terms of µOD, we first analyze the low density GFP images as follows:

**(i)** We first determine, where is the mean fluorescence intensity per pixel of a raw image and the fluorescence background which is the mean intensity per pixel in regions without cells.
**(ii)** Then we determine the number of cells,, contributing to using the following Matlab® counting algorithm that binarizes the image taking into account the inhomogeneity of the background by applying a local filter:

filename = ’raw_image’.tif’;

a=double(imread(fullfile(folder,filename)));

b=conv2(a,ones(3),’same’); c=imtophat(b,ones(20)); d=double(c>72); e=imopen(d,ones(4)); f=imreconstruct(e,d);

%N determination taking into account the systematic 20

% underestimation due to the

%presence of aggregates in images exhibiting approx. in between 200 and 1000 cells N=size(regionprops(bwconncomp(f)),1)*1.2;

A typical example is shown in Figure S5 a and b. A few images randomly picked are “manually” counted using imageJ counter tool and compared with the automatized determinations. The manually and automatized counting results are in good agreement with an average relative error of 3.5%, confirming the accuracy of the counting routine (Table S2)

**(iii)** Finally, given the linearity between and the µOD signal (Figure S6), we determine the conversion factor that relates the GFP with the µOD by means of the following formula:

Then, knowing that the number of cell *N* depends on µOD through the parameter *p* defined as: we can combine the previous two equations to define the *p* function as:

This allows to measure the value of p which allows for the determination of N(t) as a function of μOD.

We stress the fact that the proportionality factors are linked to the acquisition conditions such as the excitation intensity, acquisition time or objective numerical aperture, and must be redetermined for any change in the experimental settings.

#### PI fluorescence images analysis for cell death evaluation

To derive the number of dead cells from PI fluorescence images, we apply the same cell counting algorithm to red fluorescence images which enables accurate delineation of the cells and provide dead cells fluorescence, with the mean fluorescence intensity per pixel of a raw image (Figure S7a) and the red fluorescence background which is the mean intensity per pixel in the absence of dead cells (before irradiation). After background subtraction (Figure S7b), the binarization (Fig. S7c) allows to determine, the number of dead cells and derive, the cell fluorescence weight:

That will be used to deduce from any image of irradiated biofilm supplied with PI and recorded in the same conditions.

### Supplementary Information VI: Propidium iodide innocuity test

In order to check the innocuity of propidium iodide to PA01 PA01-GFP cells, a wide range of PI concentrations was tested against planktonic cells.

PA01 and PA01-GFP cultures were diluted in PI-enriched M9CA media to an OD of 0.05 and with PI concentration resulting in the following values: 0, 0.1875, 0.375, 0.750, 1.5, 3, 6, 12, 24, 48, 96 µM. The experiments were performed in 3 replicates per strain and concentration, while 2 controls, without bacteria but with added PI, were prepared and worked as blanks and sterility controls.

The growth was followed using the plate reader Tecan Infinite 200 Pro (Tecan, Switzerland), which kept the plate at 37 °C (Figure S8). Every 10 minutes, the plate was stirred (orbital shaking, 2 mm) and absorbance lectures were made at 600 nm. Blank absorbance was subtracted from the raw data.

When comparing the growing curves and the growth at 24 hours, both the final OD and the growth rate were not affected by the presence of PI into the media even at the highest PI concentration. No negative effect was found with the tested PI conditions against the control conditions by employing the one-way ANOVA, multiple comparisons test p > 0.05 for the growth rate and final OD at 24 hours.

**Supplementary Material References**

## Author Contributions

GI: methodology, software, formal analysis, writing - review and editing. NM-C: validation, investigation – review and editing, visualization. TP: methodology, validation. GR: conceptualization, resources, writing – review and editing, supervision, project administration, funding acquisition. NH: conceptualization, formal analysis, validation, resources, writing – original draft, writing – review and editing, supervision, project administration, funding acquisition.

## Legends to Movies S1 and S2

Movie S1: Biofilm growth without irradiation.

Typical movie from transmitted light time-lapse microscopy of a non-irradiated PAO1 biofilm growing in a PDMS-glass channel (1mmx0.25mmx30mm) supplied with M9CA medium. Focus is on top surface of the channel using 40x objective (NA 0.6). Acquisition frequency is 6 images per hour.

Movie S2: same as movie S1 for a different position.

Movie S3: PI fluorescence time-lapse microscopy of PAO1 biofilms in the presence of 3µM PI irradiated at 2h of biofilm growth. Acquisition frequency is 12 images per hour and acquisition time is 50ms.

Movie S4: same as movie S3 for a different position.

---

**Figure S1:**
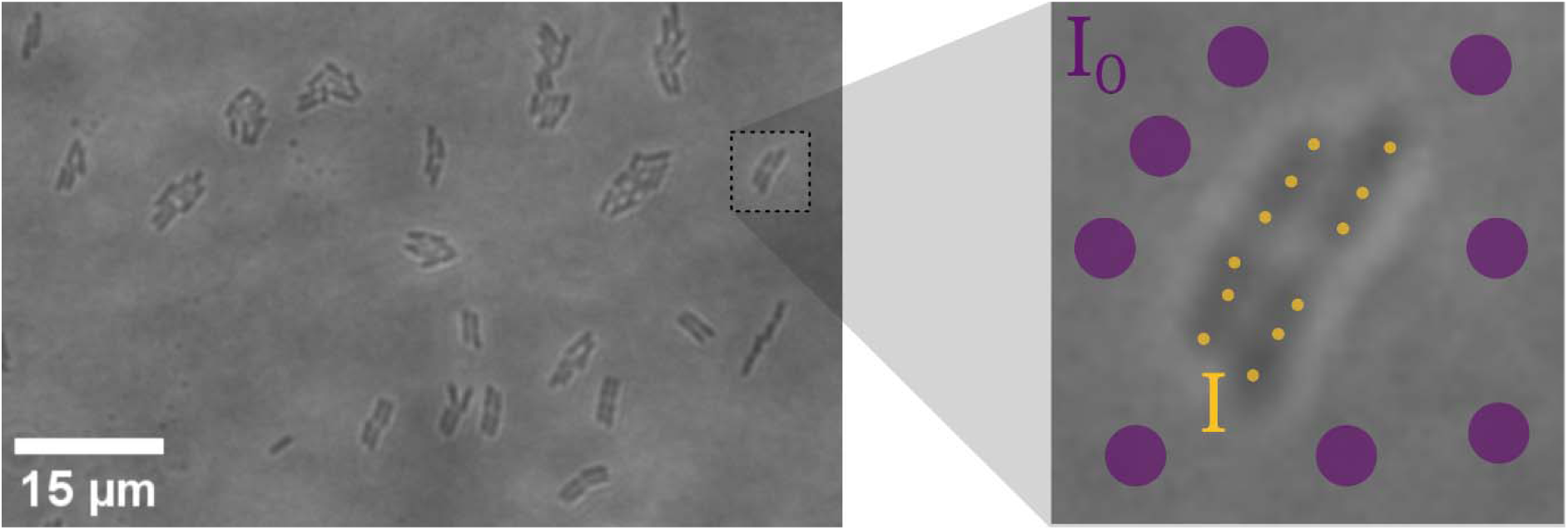
Cellular absorption coefficient. Transmitted light images of exponentially growing PA01 cells (left). Inset is represented in the right image (see text for details).

**Figure S2:**
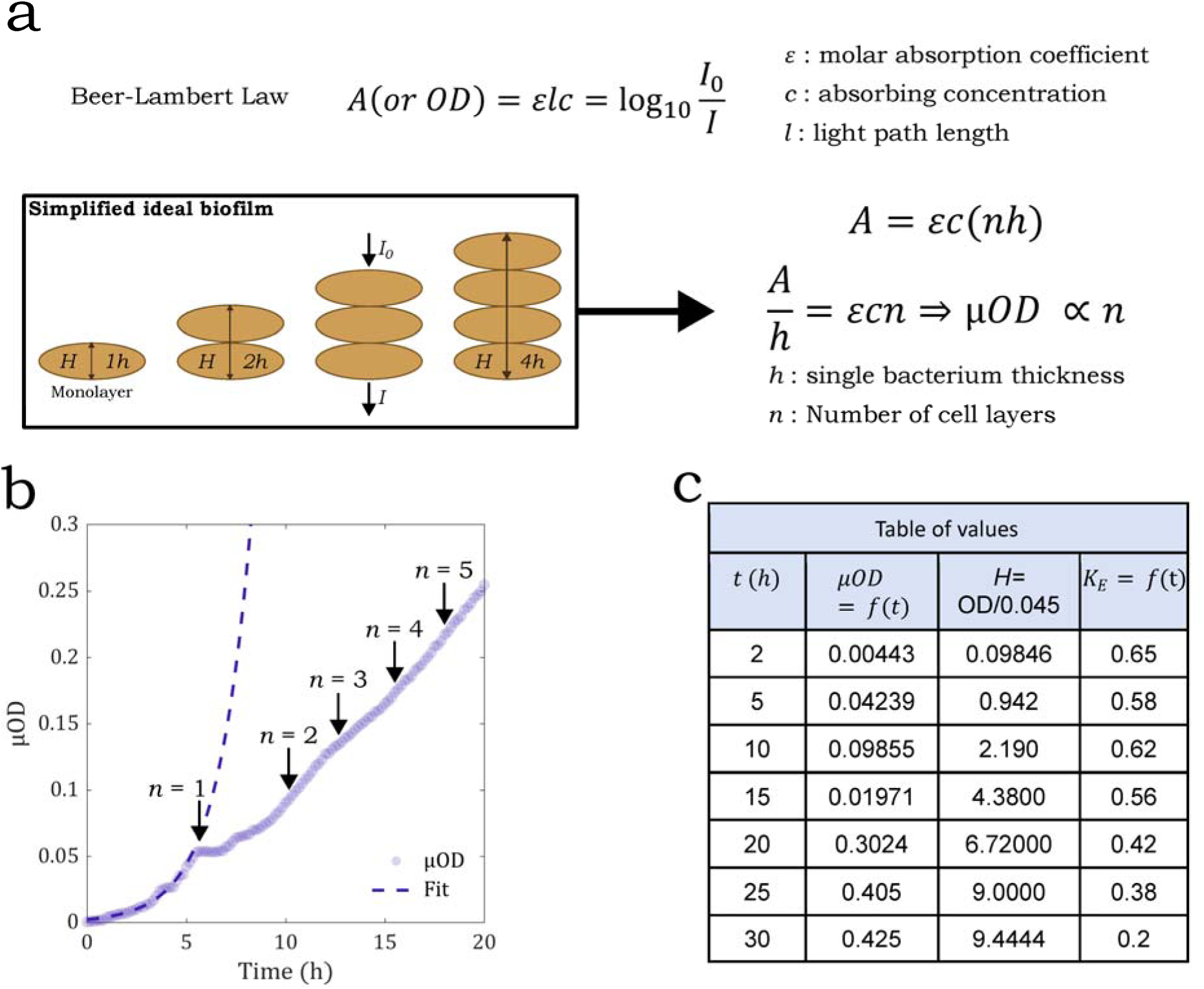
**Conversion features**. The Beer-Lambert law can be used to relate the *µOD* with the number of layers *n* as explained in panel (a). Knowing the monolayer stage (n=1) corresponds to the kink of the biofilm growth curve, we derive the conversion factor that links *µ*OD and, resulting to be 0.045±0.005 *µ*OD units per biofilm layer (average of three biological replicates) and allowing to relate *µOD, t* and *n* as marked by the black arrows (c) Then the correspondence between the *K_E_* value at the specific time and the corresponding cell layer thickness is transitively established as shown in panel c, providing the data to plot curve (Figure 6e of the main text).

**Figure S3:**
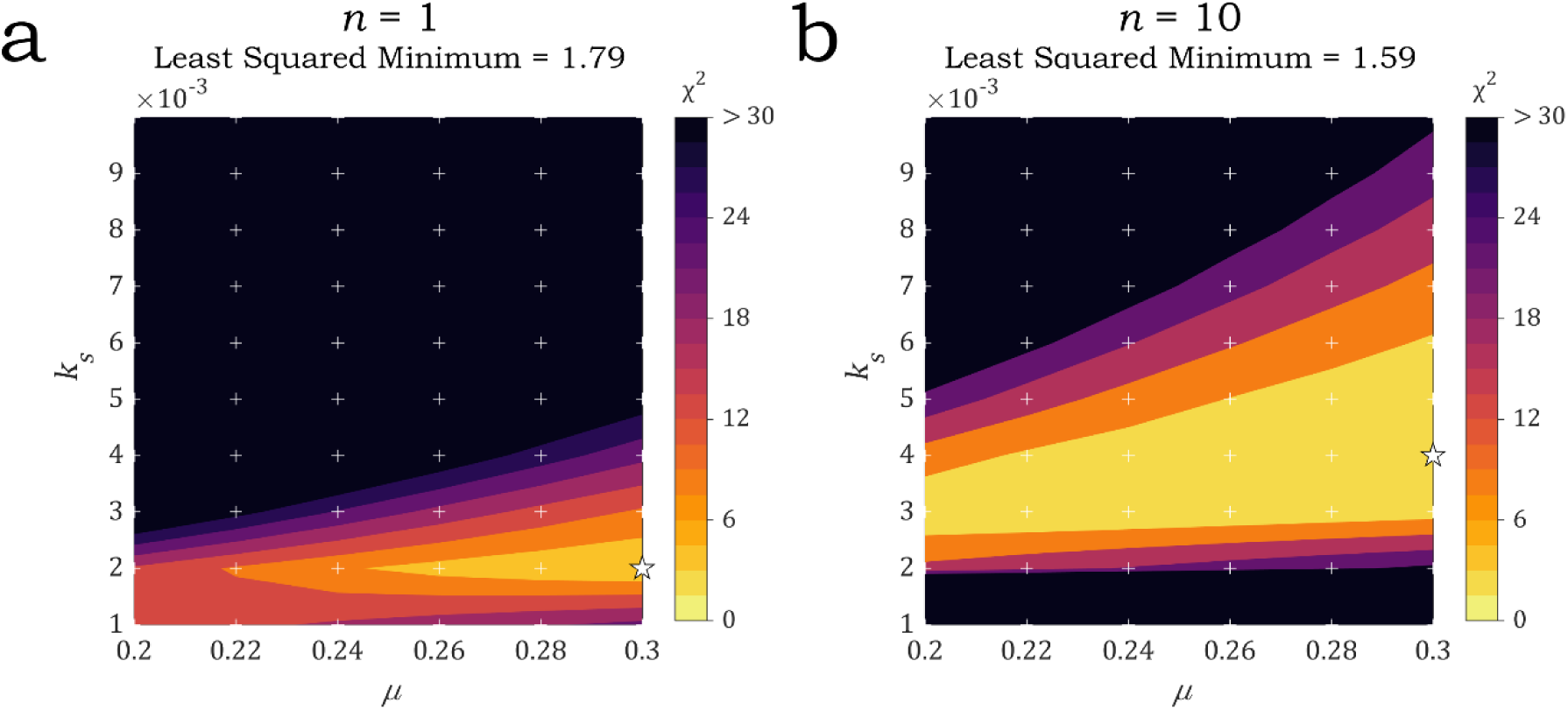
results for a light dose of 2007 J/cm², shown for = 1 (a) and = 10 (b). The other parameters vary according to the axes in the figure. The black star indicates the parameter set corresponding to the minimum Chi-squared value. The white crosses represent the parameter values used to evaluate the Chi-Squared.

**Figure S4.**
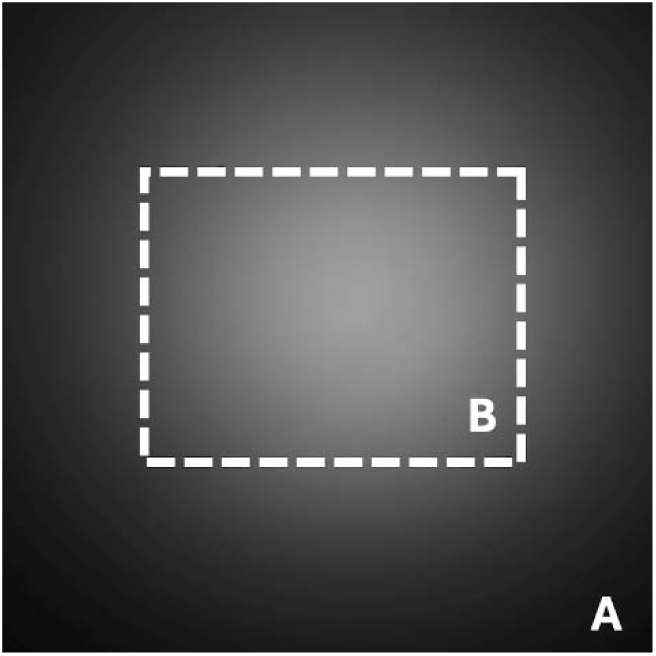
Fluorescent image corresponding to 405 nm excitation (A, 370 *µ*m x 370 *µ*m); the dotted line rectangle indicates the biofilm imaged region (B, 215 *µ*m x 165 *µ*m).

**Figure S5.**
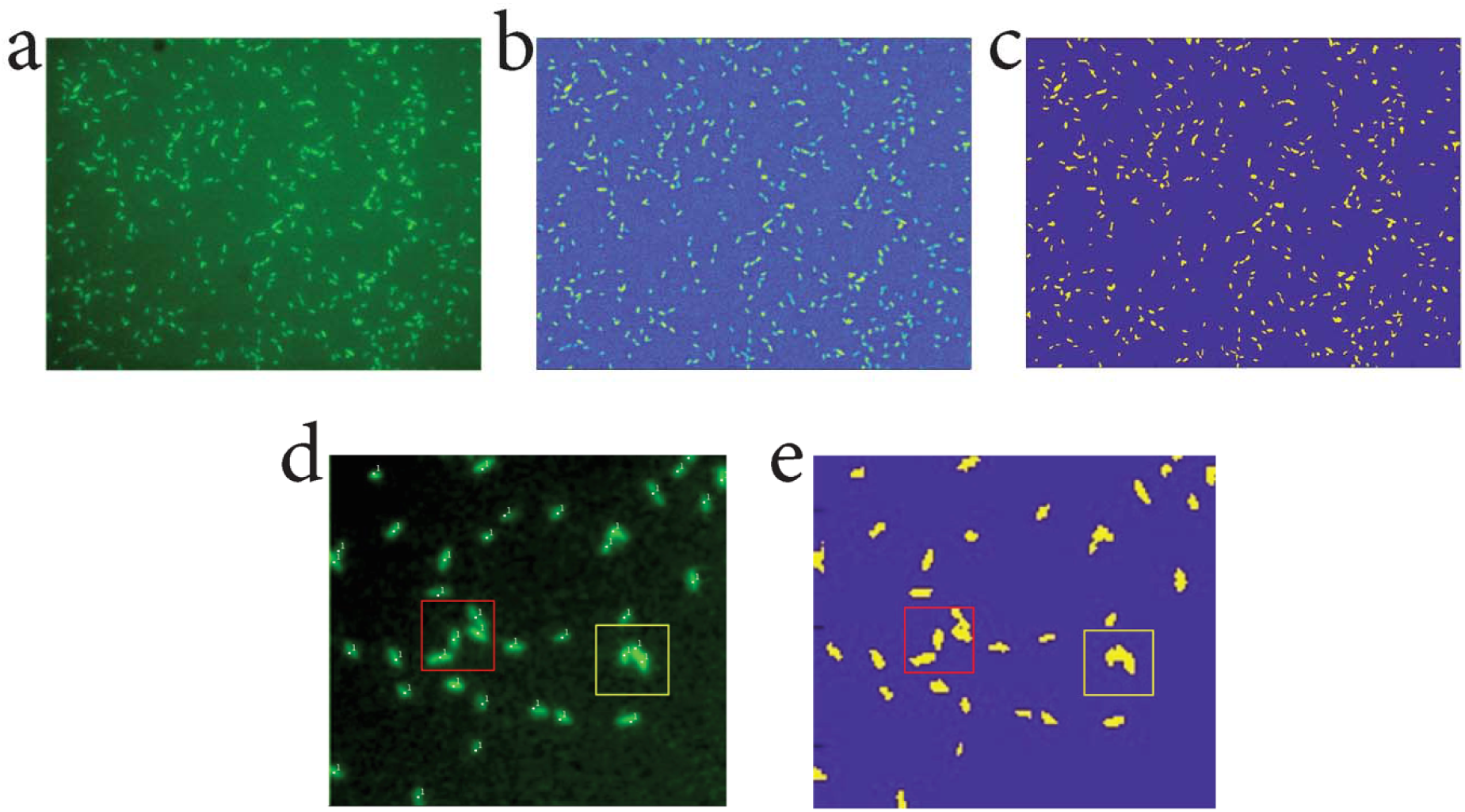
**Cell counting from GFP fluorescence images**. Images raw (a), substracted background (b) and binarized (c) are shown from left to right, respectively. Image details from the raw (d) and binarized images (e), focus on the presence of doublets or at most triplets responsible for the 20% underestimation of the automatized counting (see Table S3). Next, in the GFP linear regime, i.e. t < 4.5 hours, we calculate the GFP unit cell fluorescence weight, as follows: where includes the in-plane emission of the cells but also the contributions of the scattered and reflected light which increase linearly with the number of cells as can be shown by analyzing the mean fluorescence of empty spaces in all the images in which cell clusters can be delineated. These contributions can be viewed as a cell-dependent additional background associated with each cell.

**Figure S6.**
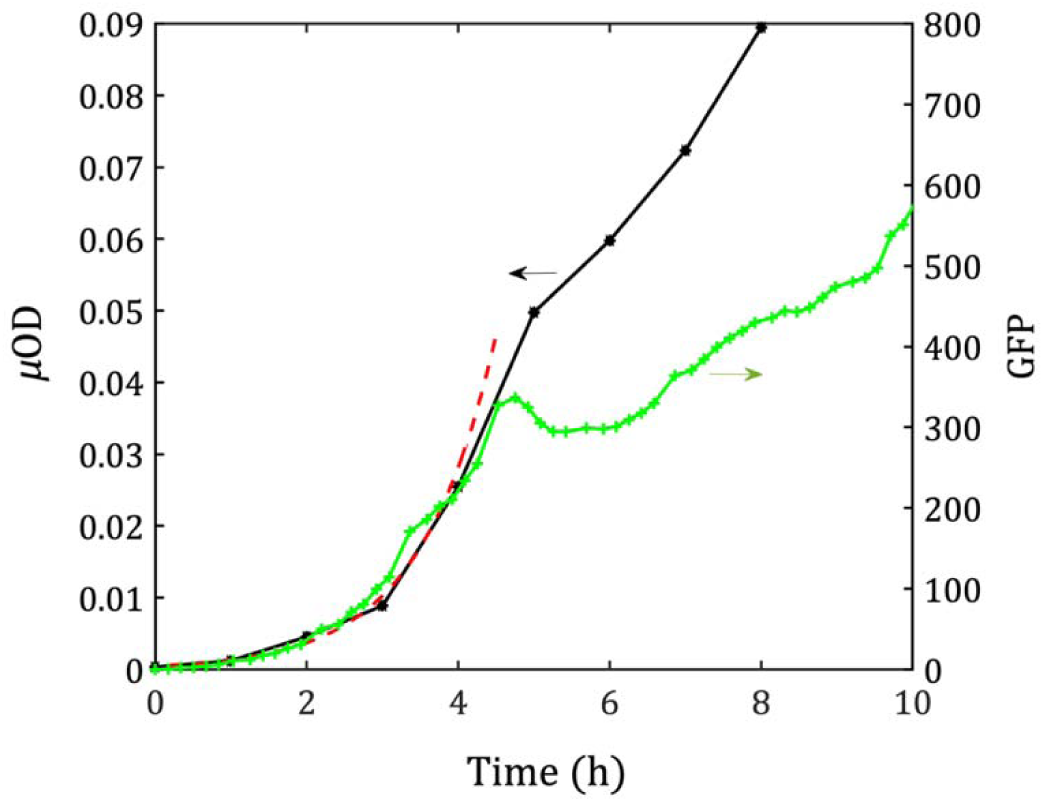
**Initial GFP vs µOD collinearity**. GFP (green curve) exhibits collinearity with the µOD curve (blue curve) up to approximately- 4.5 hours. In this time range, both signals increase according to an exponential function of the type with = 1.05 h^-1^ (purple dashed line).

**Figure S7.**
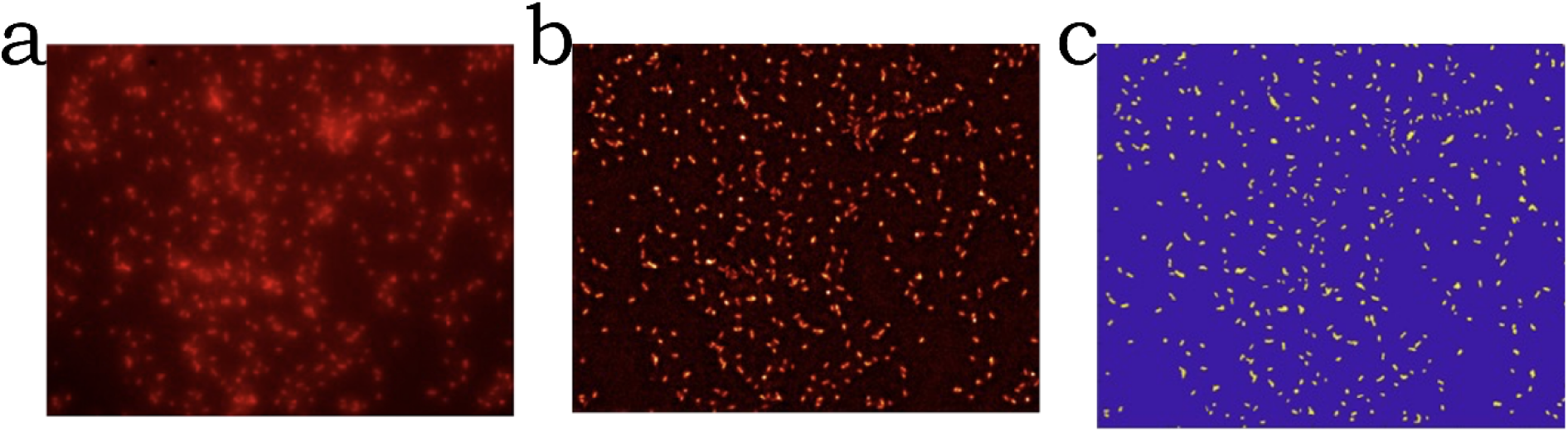
**PI fluorescence image analysis**. Typical analysis of a red fluorescence image of a biofilm supplied with PI 3 µM and irradiated after 2 hours of growth. Raw image (a) is taken at PI fluorescence intensity plateau which establishes within the few hours following the irradiation as shown in the main text. 1. (b) is the background subtracted image and (c) is the binarized image (here the image corresponds to).

**Figure S8.**
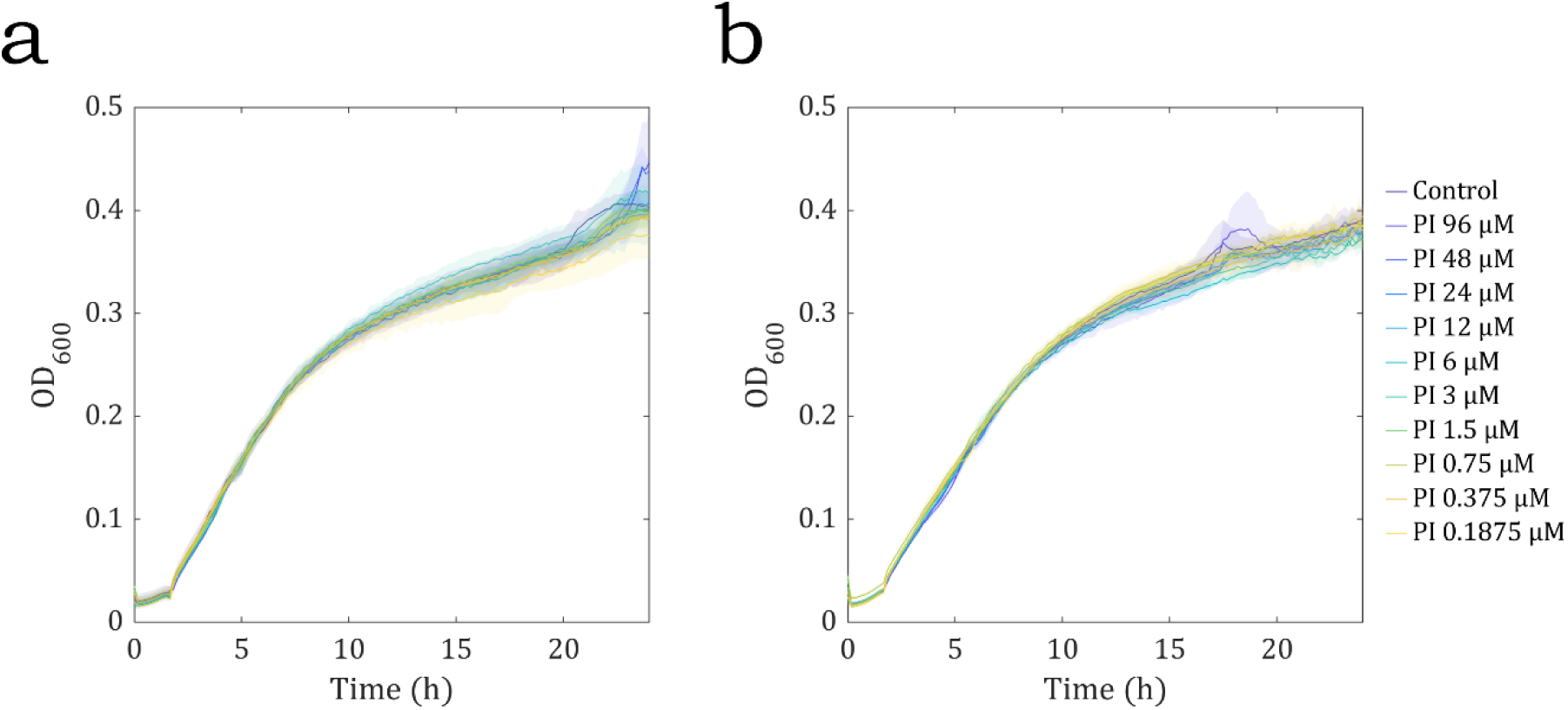
**PI toxicity**. Optical density at 600 nm (OD_600_) of PA01 (A) and PA01-GFP (B) in the presence of different PI concentrations during 24 hours.

**Table S1:** Microscopy measurement of the absorption coefficient of one cell height.

| Field of view | $I_0$ (SD)<sup>a</sup> | $I$ (SD) | μ<sup>b</sup> |
| --- | --- | --- | --- |
| #1 | 812 (39) | 673 (32) | 0.18 ±0.02 |
| #2 | 340 (12) | 286 (6) | 0.17 ±0.01 |
| #3 | 113 (4) | 86 (6) | 0.27±0.1 |
| #4 | 707 (23) | 595 (23) | 0.17 ±0.012 |
| #5 | 676 (33) | 535 (39) | 0.23 ±0.018 |
| #6 | 1607 (26) | 1279 (59) | 0.23±0.062 |
| #7 | 958 (35) | 777 (24) | 0.21±0.066 |
| #8 | 702 (29) | 510 (36) | 0.32±0.12 |
| #9 | 566(23) | 456 (26) | 0.22±0.1 |
| #10 | 2589(67) | 2084(144) | 0.22±0.093 |
|  |  |  | **0.22±0.06** |

<sup>a</sup>The measurement were performed at different intensities of incident light to rule out possible bias due to illumination.
<sup>b</sup>SD on $\mu$ is calculated from the sum of the relative SDs on $I_0$ and $I$ .

**Table S2:** Counting algorithm validation. Images from independent positions and channels are randomly picked up and counted both manually and automatically

| Algorithm count | Ground truth (Manual count) | Relative difference |
| --- | --- | --- |
| 987 | 950 | 3,8% |
| 768 | 772 | 0,5% |
| 531 | 530 | 0,1% |
| 586 | 547 | 7% |
| 518 | 494 | 4,9% |
| 979 | 1010 | 3,0% |
| 389 | 378 | 2.9% |
| 887 | 841 | 5.4% |
|  | mean | 3.5% |

**Table S3:**
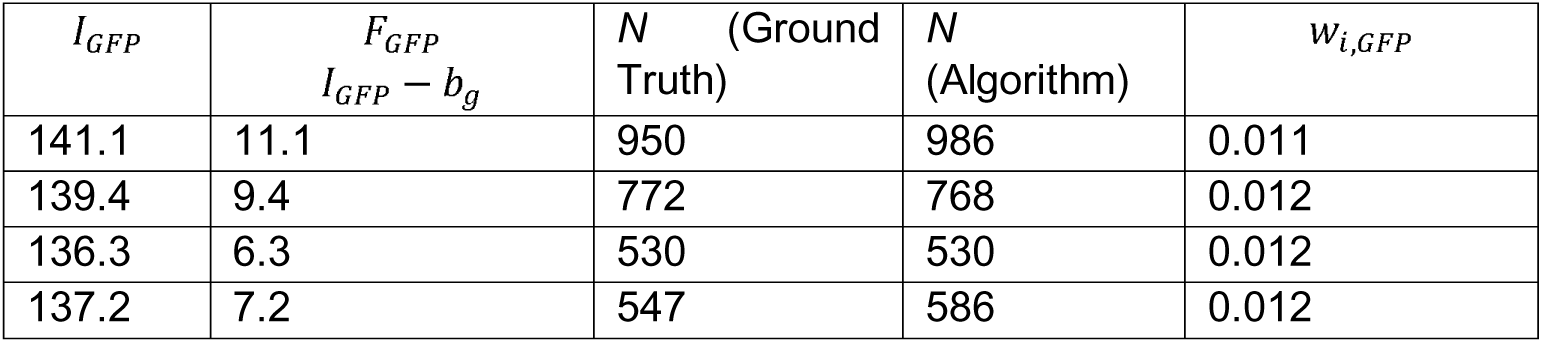
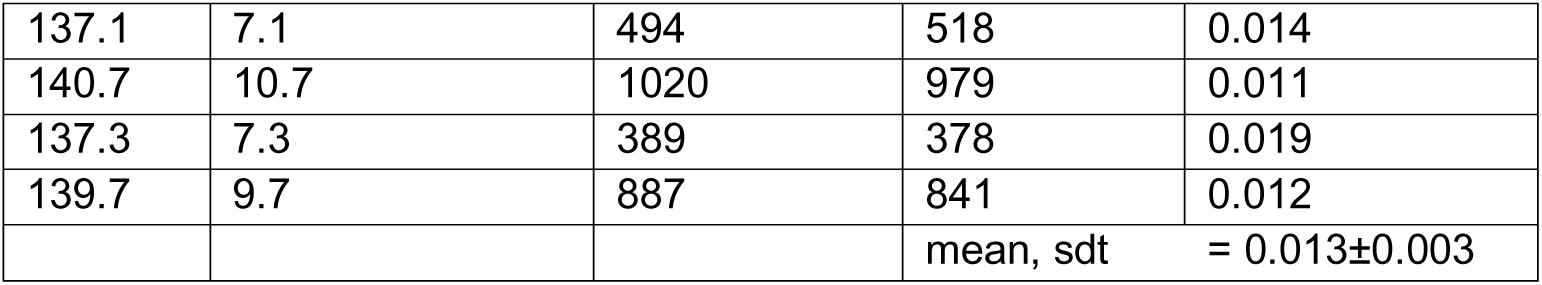
Cell fluorescent weight. Eight determinations on distinct positions in one experimental channel providing a value of 0.013±0.003 (obtained by using the automatized evaluation of N).

